# A dominant frontoparietal beta oscillatory brain state in the days after psilocybin and 5-MeO-DMT

**DOI:** 10.64898/2026.08.23.746502

**Authors:** Chloe. L West, Bradley T. Baker, Annabel Duran, Samen Nadeem, Vince C. Calhoun, Neil Van Leeuwen, Jordan P. Hamm

## Abstract

**Background:** Serotonergic psychedelics show promise for treating psychiatric disorders, with symptom improvements lasting for weeks after a single dose. Clarifying the neural basis of these effects would benefit from an identification of empirical biomarkers of such lasting shifts in brain function. Resting state EEG offers a rapid (<5 minute), low-cost window into functional brain networks. However, connectivity is not static, but cycles between recurring semi-stable patterns that vary across frequency bands. Here we employed a dynamic function connectivity (dFC) framework to identify frequency-specific connectivity states and examine how they change in the weeks following psychedelic use.

**Methods:** We collected resting-state EEG from individuals who had used one of two serotonergic psychedelic subclasses within the prior three weeks, psilocybin/LSD (typical; n=14) or 5-MeO-DMT (atypical; n=12), and age- and sex-matched controls (n=16). Frequency-band-specific spatial connectivity states (phase-lag index) were estimated across the full sample (5 per band). Groups were compared on proportion and dwell-time (per state) and state-to-state transitions.

**Results:** A right frontoparietally-distributed beta synchrony state was dominant after both typical and atypical psychedelics use (proportion/dwell-time). This effect correlated with the number of days since using psychedelics. A globally-distributed theta-band state was prominent in recent users of typical psychedelics but occurred less often in atypical users. In contrast, neural entropy (Lempel-Ziv complexity; known to increase acutely during psychedelic dosing) was not altered in recent users of either subclass.

**Conclusion:** These results reveal a beta-band signature of altered neural dynamics in the week following a psychedelic dose, consistent with a relaxation of brain network hierarchy after psychedelics.

## 1 Introduction

The acute neural effects of psychedelics are increasingly well-characterized, yet the lasting changes in brain network dynamics that follow the psychedelic experience remain poorly understood, particularly across pharmacologically distinct psychedelic compounds. Serotonergic psychedelics such as psilocybin, LSD, and DMT have emerged as promising treatments for psychiatric disorders such as major depressive disorder and post-traumatic stress disorder, with therapeutic effects persisting weeks after a single dose^1^ An understanding of the persisting changes in brain networks that accompany a psychedelic dose in this post-acute period is crucial for understanding the core mechanisms underlying therapeutic effects. Further, knowledge of which changes are common vs which are distinct between different classes of psychedelics would add additional insight, especially as details are brought to light through clinical studies.

Functional magnetic resonance imaging (fMRI) studies have provided evidence that serotonergic psychedelics elicit acute increases in global functional connectivity, i.e. widespread synchronized activity across distributed brain regions^2,3^. This widespread connectivity is associated with the disruption of hierarchical and modular brain activity structures, resulting in a more integrated and less differentiated network of neural communication^3,4^. In the post-acute period, there is more desynchronization at the global level, but the desegregating effect brain-wide networks persist last for weeks after a dose^5,6^. Although such findings are informative, Blood oxygenation-level dependent (BOLD) signal connectivity represents a low-pass filtered (<1Hz) estimate of hemodynamic correlations, which is complementary but indirectly related to changes in the principal time domains that neurons operate (1-200Hz).

Electroencephalography (EEG) studies, despite having less spatial resolution and sensitivity to areas deeper than the neocortex, provide a detailed picture of mesoscale cortical network dynamics with precise temporal resolution. Most past resting-state EEG studies have focused on the acute dosing period (1-4 hours after psilocybin, or 1 to 12 hours after LSD) and typically reveal a desynchronization of alpha-band oscillations and increased signal complexity^7^. Whether these changes persist in the post-acute period is unknown, yet is crucial for building a more comprehensive understanding of the clinical effects of these compounds. Notably, animal studies document spine growth in cortico-cortical projecting neurons which do not peak until at least 24hrs after the dose but last for months^8,9^. Given that EEG is sensitive to cortical networks, one would expect that, if human brain networks exhibit similar synaptic changes to rodent studies, there should be some persisting EEG signature of psychedelic induced plasticity.

Another gap in the literature comes from the fact that past fMRI and EEG work has used static connectivity measures. Static connectivity measures, which assume stable connectivity patterns over time, fail to capture the dynamic nature of brain network activity^10–12^. This limitation is particularly significant for studying the effects of psychedelics. Psychedelic states are characterized by rapidly changing and highly variable brain activity patterns^13–15^, and more sensitive measures that account for temporal variability are needed. And – considering the afterglow effects of psychedelics^16^ along with the finding that certain psychedelics (e.g. LSD and 5-MeO-DMT) often involve sudden reactivations of the psychedelic experience well-after the acute dose (i.e., flashbacks)^17^, changes in network dynamics may extend beyond the acute period. Dynamic functional connectivity (dFC) is an alternative metric to static connectivity that captures temporal fluctuations in functional connectivity patterns, isolating distinct “states”, each characterized by unique patterns of functional connectivity^10^. States tend to be quite consistent over subjects, and as such, the number of transitions between and relative time spent within particular states can be quantified and compared^11,12,18^. dFC analysis is particularly advantageous for electroencephalography (EEG) data, capitalizing on EEG’s excellent temporal resolution to provide frequency-specific characterizations of connectivity states^11^.

Additionally, past work using EEG to study acute psychedelic states (i.e. immediately after a dose) has reliably found a temporary increase in neural entropy, reflecting an increase in signal complexity and suggesting a breakdown of segregated network activity structures^19,20^. These effects have been proposed as biomarker representative of a key therapeutic mechanism^19–22^, however, determining whether entropy remains altered after the acute psychedelic state, when basic perception mostly returns to baseline but therapeutic effects remain, is key for evaluating this proposition.

While ‘typical’ serotonergic psychedelics such as psilocybin and Lysergic acid diethylamide (LSD) primarily exert their phenomenological^23,24^ and much of their neuroplastic effects through 5-HT_2A_ receptor agonism^9,25–27^, there are other serotonergic psychedelics that produce an intense phenomenological psychedelic experiences, despite having divergent receptor binding profiles – namely, 5-methoxy-N,N-dimethyltryptamine (5-MeO-DMT). This “atypical” serotonergic psychedelic, 5-MeO-DMT, displays a distinct pharmacological profile with a 100-fold higher affinity for the 5-HT_1A_ receptor over the 5-HT_2A_ receptor^28^, and, notably, 5-HT_1A_ receptor agonism often antagonizes 5-HT_2A_ receptor activity^29^. Compared to typical psychedelics, the 5-MeO-DMT psychoactive experience is much less visually driven and can even include “white-out experiences” reminiscent of amnesia or a void-like state^28^. Nevertheless, 5-MeO-DMT exerts comparable therapeutic^28^ and neuroplastic effects^30^ to typical serotonergic psychedelics^31^. Furthermore, the short duration of action characteristic of 5-MeO-DMT (15-30 minutes^32^) makes it an attractive option in therapeutic settings compared to longer-lasting psychedelics like LSD and psilocybin^24^. Comparing typical (e.g., psilocybin) and atypical serotonergic psychedelics offers a means to test whether post-acute connectivity changes are 5-HT_2A_ receptor dependent^33^ and to characterize whether distinct pharmacological mechanisms engage different pathways to therapeutic outcomes.

This study analyzed dynamic functional connectivity patterns in EEG recordings gathered from three groups: control participants, individuals who have recently used typical psychedelics (psilocybin or LSD), and individuals who have recently used the atypical psychedelic 5-MeO-DMT (< 3 weeks). We utilized the weighted phase lag index (wPLI) to assess both full spectral analysis and specific frequency bands of dFC^34^. In particular, theta and gamma oscillations are known to mediate feedforward processing in primate neocortex ^35–37^, while beta oscillations serve broader roles in inter-regional communication^36,38^, including feed-back modulation from higher to lower brain regions. Collectively, these properties outlined our overarching expectation that the post-acute period is characterized by a structured reorganization of feedforward and feedback dynamics, enhanced bottom-up processing and disrupted top-down control, that persist beyond the acute period. More broadly, we hypothesized that the typical psychedelic group would spend a greater proportion of time in globally hyper-connected states compared to controls, consistent with documented increases in global functional connectivity following typical psychedelic use^2–4^. We assessed three measures of dFC: proportion of time spent in each state, mean dwell time in each state, and the number of transitions between states.

## 2 Methods

*Ethical Approval*: All experimental procedures were approved by the Georgia State University Institutional Review Board (IRB) and were carried out in accordance with their guidelines. Participants were recruited through advertisements posted on university bulletin boards, social media platforms, and in local psychedelic community groups. Eligibility criteria included adults aged 18 or older with no history of autism spectrum disorders, bipolar disorder, schizophrenia, current major depressive episodes, or substance abuse, excluding nicotine. Participants filled out a detailed questionnaire to ensure suitability for study participation. Participants provided informed consent and were compensated $30 for their participation in the study.

The study cohort comprised 42 individuals, divided into a control group, a typical serotonergic psychedelic group, and an atypical serotonergic psychedelic group. The typical serotonergic psychedelic group included 13 participants (6 male, 7 female) who had used psilocybin and 1 (male) who had used LSD within 30 days prior to their participation in this study. The atypical serotonergic psychedelic group comprised 12 participants (5 male, 7 female) who used inhaled Incilius alvarius toad toxin. 5-MeO-DMT is considered to be the primary compound present that is responsible for the toxin’s psychoactive effects. The control group consisted of 16 age and sex matched participants (6 male, 10 female), who had no recent (within 365+ days) history of psychedelic use. See table 1 for detailed demographic data. Dosage information for psilocybin, LSD, or 5-MEO-DMT was unattainable, as potency of the drugs used could not be ascertained directly, but participants were pre-screened to exclude those who only took sub-perceptual microdoses. All doses reported were judged as moderate to high.

**Table 1:**
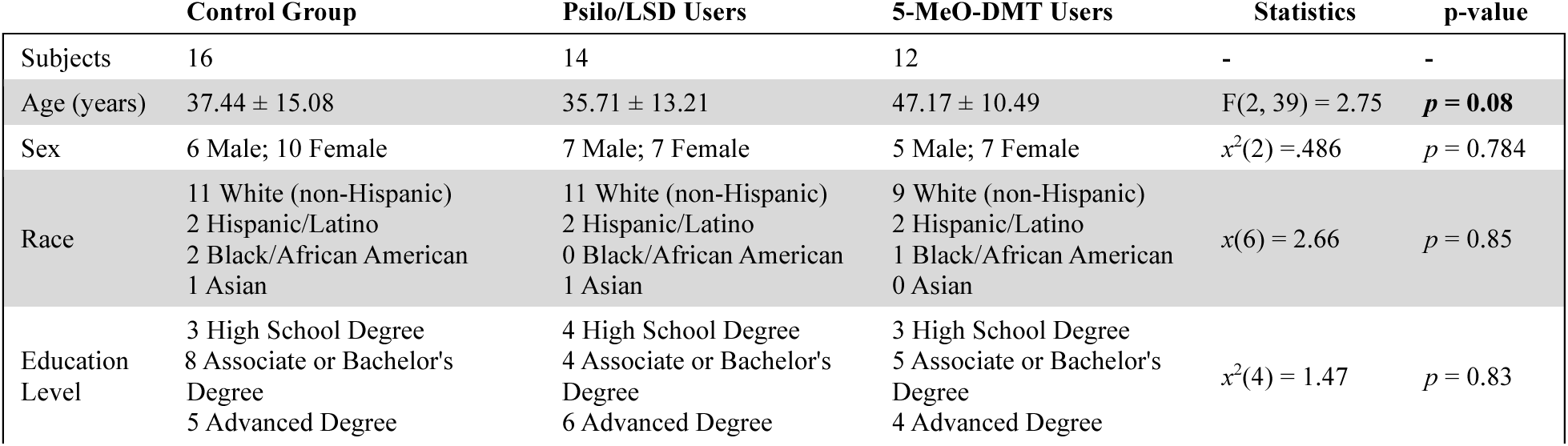
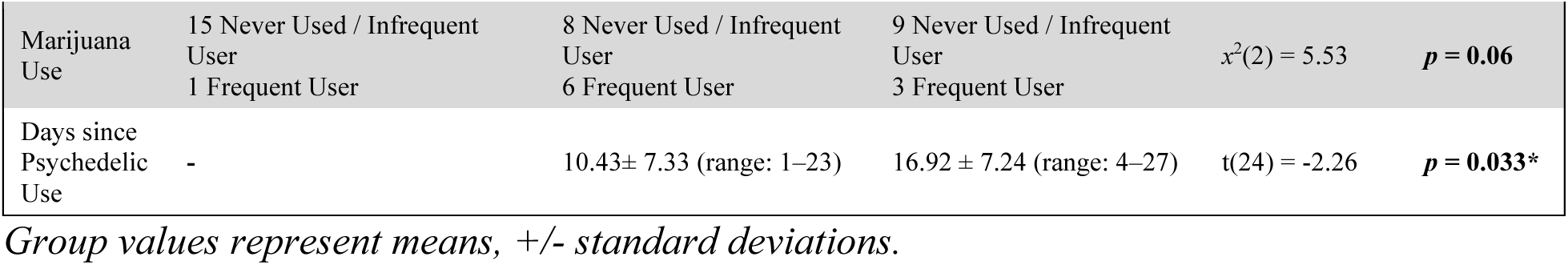
Participant demographic and substance use characteristics.

### 2.1 Resting-State EEG recordings and processing

EEG data were recorded in a darkened room with moderate temperature, moderate humidity, good ventilation, and electromagnetic insulation. Participants were seated approximately 40 inches away from an LCD monitor (19-27 inches, 60Hz refresh rate). Participants were instructed to keep their eyes open stare at a blank blacked-out screen.

EEG recordings and data processing were conducted using a 32-channel BioSemi ActiveTwo EEG system. Participants were fitted with a BioSemi 32-channel cap, arranged according to the 10-20 system for electrode placement. The EEG data were recorded with a sampling rate of 500 Hz, using a band-pass filter of 0.16 Hz and a lowpass filter at 200 Hz. Reference electrodes were placed 1 cm to the right of Cz (later average referenced) and the ground electrode was positioned at 1 cm to the left of Cz. Independent component analysis (ICA) was used to remove ocular and muscular artifacts, followed by manual visual inspection to ensure the accuracy of artifact removal (BESA software). The cleaned EEG data were then segmented into 1-second epochs. Average power spectra for each electrode for each group is plotted in figure S1. This was not compared between groups and is presented for visualization purposes (see rationale in 2.2 below).

### 2.2 Weighted Phase Lag Index

Let X and Y be time series with length T. The wPLI^34^ measures how differences in the phase angle between X and Y tend to be positively or negatively distributed along an imaginary axis of the complex plane. Standard PLI is defined as:

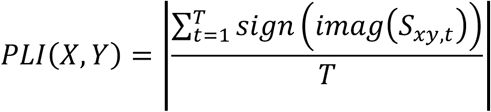

where S_xy_ is the cross-spectral density of X and Y at time T. While PLI is insensitive to zero-lag interactions, noise and other confounds can be introduced via volume conduction.

The wPLI, defined as:

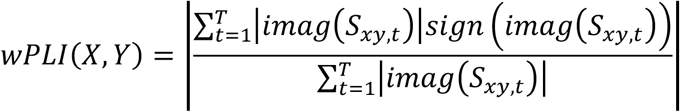

addresses the limitation of PLI by scaling phase angle differences by their distance from the real line. PLI and wPLI both utilize only the imaginary part of the cross-spectral density, and are robust to noise when compared to other FC metrics like coherence or Pearson’s correlation coefficient. For this work, we focus only on wPLI, leveraging the robustness to noise and volume conduction established in the literature. Static connectivity for each frequency band, for each group is plotted in supplemental figures S3, S5, S7, S9, and S11. We chose not to statically compare average power (Figure S2) or general wPLI connectivity matrices (Figures S3, S5, S7, S9, and S11) as these are static measures. The focus of this study was dFC. If differences in network dynamics are present in the data (e.g. more time spent in high connectivity states, but not a true shift in overall functional connectivity), it could ostensibly appear as overall changes in these static measures. Statistical comparisons of these static measures could therefore conflate changes in state occupancy or other dynamic properties with changes in overall network connectivity. We therefore focused our inferential analyses on the dynamic measures that more directly address the study hypotheses.

### 2.3 Dynamic Functional Connectivity

Dynamic functional connectivity was assessed using a sliding window approach, with windows of 5 seconds duration and 1-second overlap, following the precedent in Chen et al.^11^. Connectivity within each window was quantified using the wPLI. dFC analysis was conducted across the entire frequency spectrum (full spectral analysis) and within individual frequency bands: delta (1–4 Hz), theta (5–8 Hz), alpha (9–14 Hz), beta (15–25 Hz), and gamma (26–40 Hz).

To perform temporal segmentation, k-means clustering was performed on the wPLI connectivity windows within each band to identify distinct brain states. This clustering approach allows for a data-driven association of related connectivity windows, and while it is just one method for determination of states, it is well-established in the functional connectivity literature^18,39–41^. Different values of K between 2 and 16 were evaluated, with K=5 determined to be the optimal number according to the elbow criterion using the silhouette score^42^ and Davies-Bouldin index^43^, which evaluates the variance explained by clusters and identifies the point at which adding more clusters provides diminishing returns. K=5 was optimal for all frequency bands. For computational efficiency, the elbow criterion was evaluated on a subset of exemplar windows computed for each participant representing local maxima of the standard deviation computed over windows. Each exemplar clustering was initialized randomly 10 times and the result with the optimal tradeoff in scores was taken. The resulting states from the clustering of exemplar windows were then utilized to initialize the clustering of the full set of windows. The final cluster analyses were carried out on all data across all subjects together, separately for each frequency band. K-Means was implemented using the scikit-learn package (v0.24.2) in Python 3.9.12. Each set of states is plotted for each frequency band in Supplemental figures S4, S6, S8, S10, and S12.

Following K-Means clustering, ongoing activity during the recordings for each subject, for each frequency band, for each time-bin, was assigned to one of these 5 canonical brain states (clusters) based on a Euclidean distance criterion. Then, for each subject and for each band, we calculated proportion of time in each state, mean dwell time in each state, and number of transitions (over all states, within a frequency band). Proportion time for state *k* is measured as the total time that a participant spends within that state divided by the total number of windows in the recording. Mean dwell time is computed as the average duration that a particular state was maintained before transitioning to another state. Finally, the number of transitions quantifies the total number of transitions between different states, reflecting the stability and dynamism of the brain’s connectivity patterns. To test for group differences in proportion time and mean dwell time, we performed linear mixed effects analyses for each frequency band, aimed at identifying STATE by GROUP interaction effects, with age and sex as fixed effects. We followed-up significant interactions with pairwise t-tests for each state, using a False-discovery rate adjustment for multiple comparisons. Number of transitions was tested using a similar model, but without STATE. Results are reported in figures 1–4.

**Figure 1.**
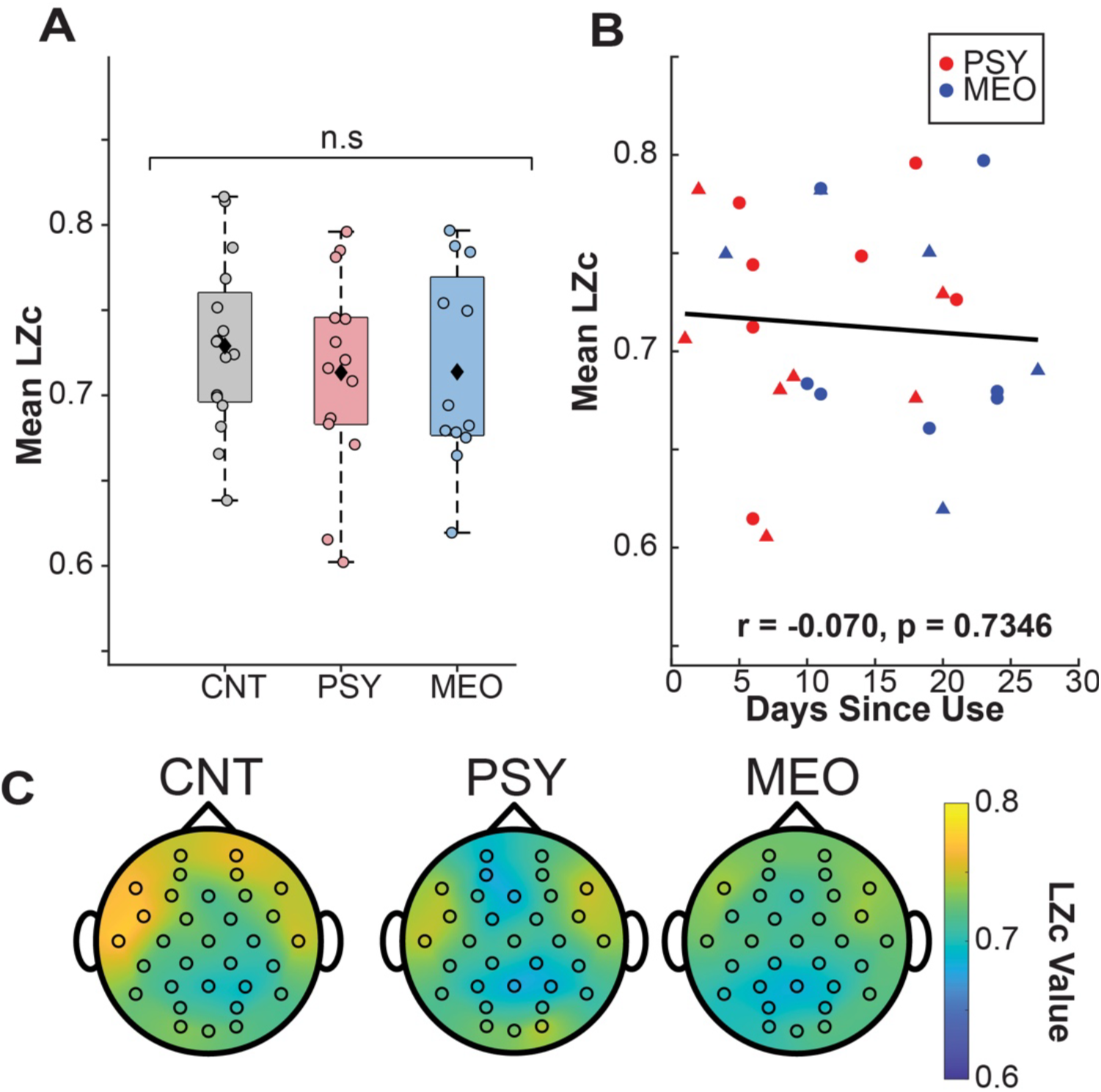
Lempel-Ziv Complexity (LZc) does not differ across groups or correlate with days since use. (A) Average LZc values across EEG channels for each subject, grouped by use group. (B) Average LZc values averaged across channels for each subject in the PSY and MEO groups, plotted agains days since use, indicating no significant correlation. Triangles are male. C) Topographical maps depicting the spatial distribution of LZc values for each group, with color intensity corresponding to LZc magnitude. No group by region (occipital, parietal, central, frontal, temporal) interactions were present. MEO = 5-MeO-DMT group; PSY = psilocybin/LSD group; CNT = control group.

### 2.4 Lempel-Ziv Complexity

Lempel-Ziv complexity (LZC) is a measure of the complexity of a time series, capturing the degree of randomness and the number of distinct patterns within the data^21,22^. LZC quantifies the degree to which a given signal can compress, with more complex signals resulting in higher LZC values^21^. Following established methods^44^, recordings were down-sampled from 500 Hz to 250 Hz using an anti-aliasing filter, and then segmented into 3-second epochs, corresponding to 750 samples per epoch. Then the data were linearly detrended and standardized by subtracting the mean and dividing by the standard deviation for each channel, ensuring that the analysis was not influenced by amplitude differences across channels or epochs. Next, the Hilbert transform was applied to each standardized epoch to obtain the amplitude envelope of the EEG signal. A binary string representation was then generated by comparing the amplitude envelope to its mean value, with samples above the mean encoded as ‘1’ and those below the mean encoded as ‘0’. To compute LZc, the binary string was shuffled to create a surrogate dataset, and LZc was calculated as the ratio of the compression length of the original binary string to that of the shuffled string, quantifying the complexity of the EEG signal with higher values indicating greater complexity. LCZ values were then calculated for each “region” (frontal, temporal, parietal, occipital, 4 total) for each subject, and subjected to Linear mixed effects analysis to examine a GROUP by REGION interaction, with SEX and AGE as fixed effects.

## 3 Results

### 3.1 Lempel-Ziv complexity not altered in weeks following typical and atypical psychedelic use

First, we decided to explore whether entropy, measured by LZc, was elevated in our groups, given previous research showing increased entropy during psychedelic experiences. Our LZc analysis did not reveal any main effects or interactions involving group (GROUP X REGION -- *F* (6,154) = 1.34, *p* = .24; GROUP – F(2,154)=1.028, p=.36; Figure 1A,C). Within the psychedelics groups, this did not correlate with days since use (r=-.057, p=.73; Figure 1B). This contrasts past findings of an increase in LZC in the acute phase after psychedelics.

### 3.2 Band-specific dynamic functional connectivity differences in theta and beta across psychedelic users and controls

Next, we analyzed differences in frequency band-specific global dynamic functional connectivity patterns. We examined the following frequency bands: delta (1-4 Hz), theta (5-8 Hz), alpha (9-14 Hz), beta (15-25 Hz), and low-gamma (26-40 Hz). For each band, we identified five distinct states (supplemental figures S4-S12). We did not observe and main effects or interactions involving GROUP for number of transitions between states (F-GROUP (2,193) = 0.807, p = .45; F-GROUP by frequencyBAND (8,193) = 0.658, p = .73).

On the other hand, groups differed in the time that they occupied specific states for theta, alpha, and beta bands. For theta, this was apparent as GROUP by STATE interactions for proportion of time spent per state (F(8,193)=4.428, p=6.10e-5) and mean dwell time (F(8,193)=4.20, p=1.14e-4). For alpha, this was apparent as a GROUP by STATE interaction only for proportion time (F(8,193)=3.002, p=.003). For beta, this was apparent as GROUP by STATE interactions for proportion of time spent per state (F(8,193)=2.272, p=.024) and mean dwell time (F(8,193)=2.100, p=.038).

We followed up these interactions by examining orthogonal contrasts (control vs (5-MeO-DMT & Psilocybin/LSD); 5-MeO-DMT vs Psilocybin/LSD) via t-tests for each state (0 through 4) and for the above 5 effects, and adjusting p-values for multiple comparisons using FDR. Consequently, three contrasts remained statistically significant and are discussed below.

#### 3.2.1 Increased time spent in high-connectivity states in the theta band (5-8Hz) after typical psychedelics

Within the theta frequency band (5–8 Hz), we identified 5 connectivity states (see supplemental figure S6), numbered 0 through 4. Groups differed in the proportion of time occupying state 0, with the two psychedelics groups strongly differing from one another (t(39)=3.27, p=.002). Specifically, the typical (psilocybin/LSD) group exhibited a high proportion of the rest period in this state while the atypical (5-MeO-DMT) group spent comparatively little time (figure 2B). Control comparisons subjects spent an intermediate amount of time in this state.

**Figure 2.**
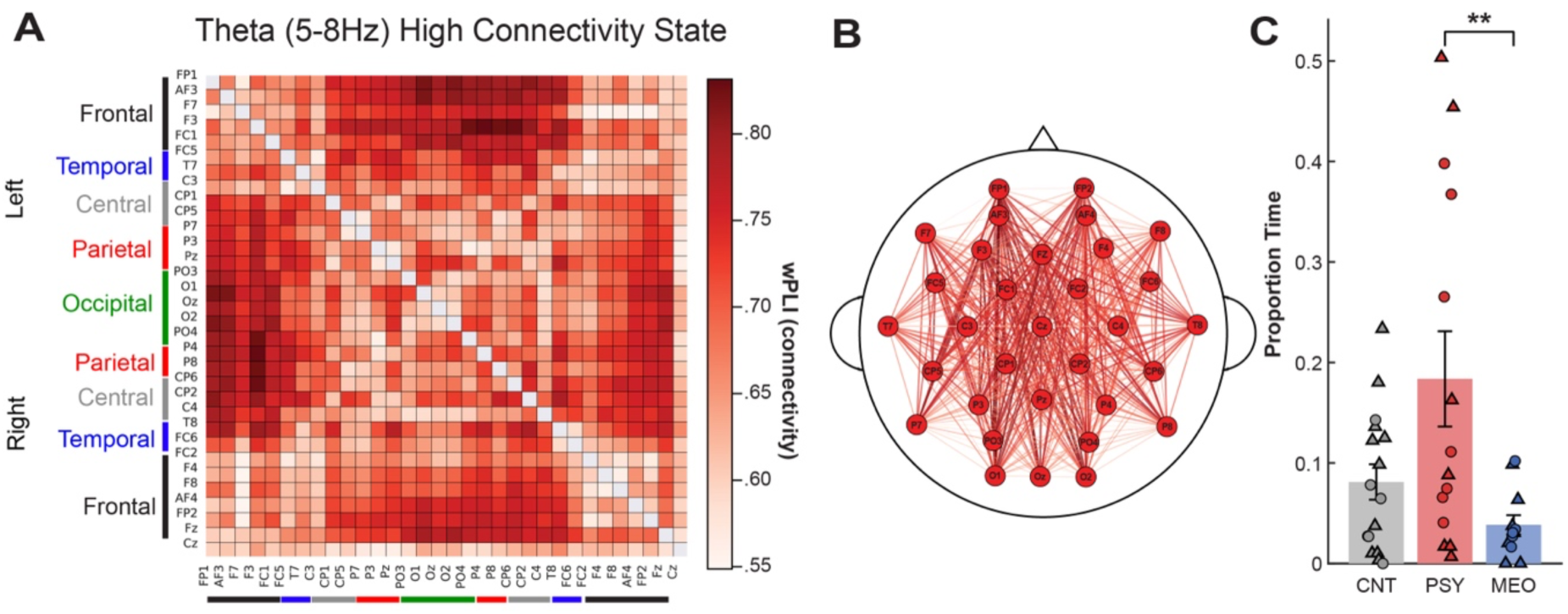
A broadly connected theta-band state differentiates psychedelic user groups. (A) Mean wPLI connectivity matrix (left) and (B) corresponding topography of connectivity (right) for theta “State 0”, depicting a global high-connectivity state characterized by elevated wPLI values across widespread electrode pairs. Line thickness and color intensity correspond to wPLI connection strength. (C) Proportion of time spent in State 0 by group, indicating group mean and standard deviations. Individual data points are overlaid. Males are triangles. \*\**p* < .01.

Visual inspection of the mean connectivity matrices revealed that State 0 represented a global high-connectivity state characterized by elevated wPLI values across widespread electrode pairs (Figure 2A). Paired-samples t-tests confirmed that theta State 0 exhibited significantly greater aggregate connectivity (M = 0.68) than all other theta states (Ms = 0.44– 0.57; all ps < 10⁻²⁴, ds = 3.95–7.13). This pattern was consistent across all pairwise comparisons, with State 0 showing the highest mean connectivity of any theta state identified by the clustering algorithm. The corresponded to dense connectivity spanning frontal, central, parietal, and occipital regions, with high wPLI values distributed broadly across the scalp.

No significant group differences were observed for proportion time or mean dwell time in States 1–4. Proportion of time spent in theta state 0 did not correlate with age (partial-r=-.162, beta=-.0018, p=.449), sex (partial-r=-.020, beta=.005, p=.962) or with since use of psychedelics (partial-r=-.31, beta=-.006, p=.13, with sex and age included in the model).

#### 3.2.2 Increased time spent in frontal beta-band connectivity states after psychedelics

K-means clustering also identified five distinct connectivity states within the beta frequency band (15–25 Hz; Supplemental figure S10). Again, groups differed in the proportion of time occupying state 0, with psychedelics groups (typical and atypical combined) exhibiting increased time spent (t(39)=3.423, p=.001) and longer average dwell times in this state (t(39)=3.255, p=.0023) than comparison subjects. Psychedelics groups did not differ from each other on these measures. Visual inspection of the mean connectivity matrices revealed that State 0 was characterized by a frontoparietal connectivity pattern, with dense connectivity among frontal, fronto-central, and fronto-parietal electrode pairs and comparatively sparser posterior connections (Fig. 3A). This pattern was also stronger in the right hemisphere. While State 0 did not exhibit the highest aggregate connectivity of the five identified states, it was distinguished by this anterior-weighted topography.

**Figure 3.**
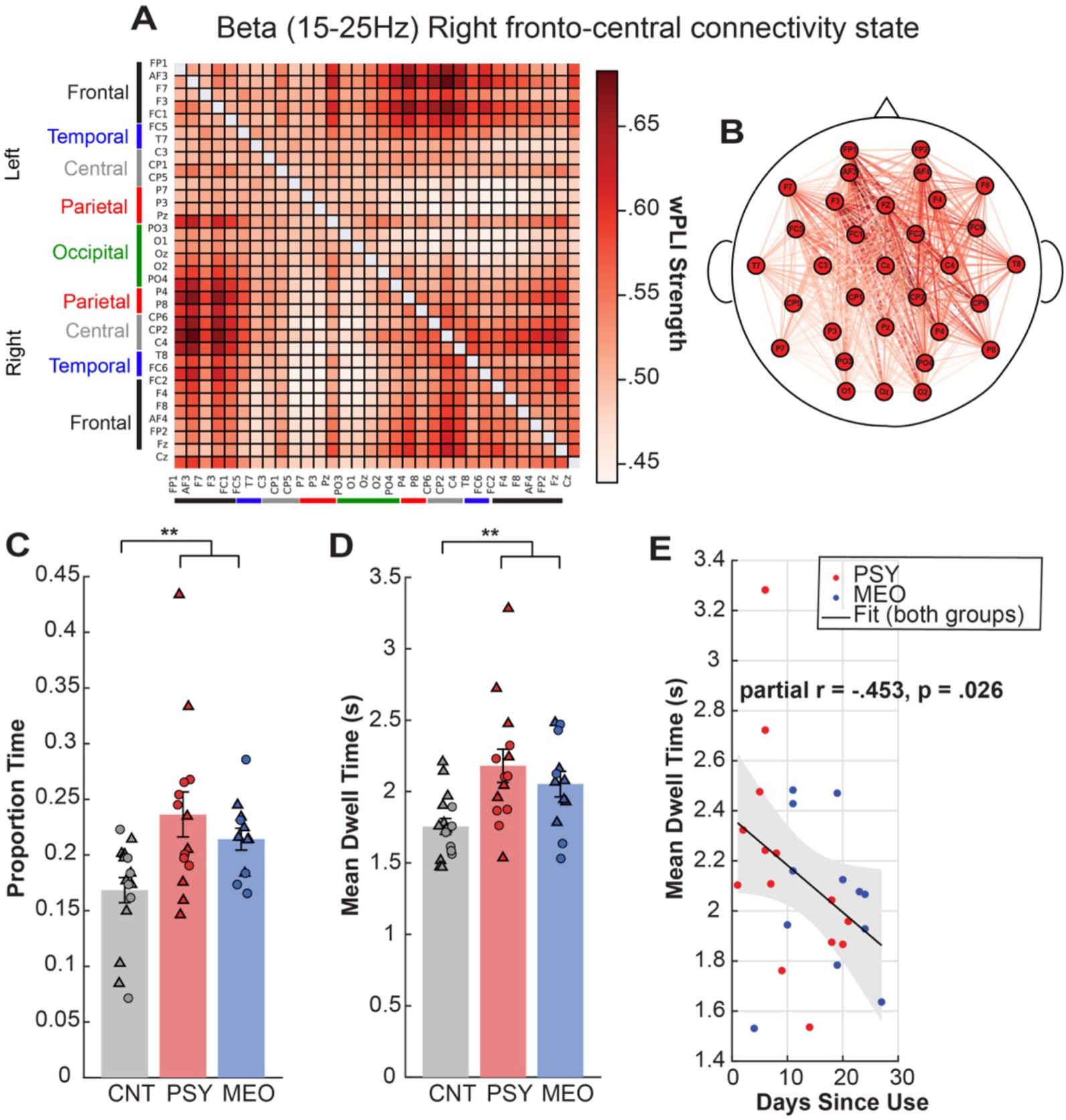
Beta (15–25 Hz) dynamic functional connectivity State 0. (A) Mean wPLI connectivity matrix and (B) corresponding connectivity topography for beta state 0, depicting a right frontoparietal-dominant state. (C) Proportion of time spent in State 0 by group, means and SEM, with individual subject scatter overlaid (male=triangle). (D) same as C but for mean dwell time in in state 0 \*\**p* < .01. (E) Scatter plot depicting the association between mean dwell time in beta State 0 and days since last psychedelic. The line represents the linear regression fit and the shaded region indicates the 95% confidence interval. *(partial r = −.453, β = −0.0226, p = .0262)*.

No significant group differences were observed for proportion time or mean dwell time in States 1–4. Proportion of time spent in beta state 0 did not correlate with age (partial-r=.336, beta=.0014, p=.108) or sex (partial-r=-.351, beta=-.0039, p=.393), but it did negatively correlate with days since use of psychedelics (partial r = −.411, β = −0.003, p = .046), returning to the levels of control participants by three weeks post dose. The same pattern was observed for mean dwell time in beta state 0; it did not correlate with age (partial-r=.222, beta=-.006, p=..298) or sex (partial-r=-.348, beta=-.237, p=.398), but it also negatively correlated with days since use of psychedelics (partial r = −.453, β = −0.0226, p = .0262), returning to the levels of control participants after 3 weeks.

## 4 Discussion

These results present the first dFC study of resting state EEG after the use of psychedelics. dFC provides a unique quantification of resting brain function that moves beyond static estimates of power or patterns of inter-electrode connectivity, which overlook changing patterns and semi-stable, dominant network states^10^. Here we identify a fronto-parietally distributed beta-synchronous brain state that becomes more dominant in individuals taking both typical and atypical serotonergic psychedelics in the first week after use, and tapers to non-user levels after that. Further, we find a specific global theta-synchronous brain state that differentiates typical vs atypical psychedelics.

Together, these findings fit with converging evidence that suggests psychedelics produce lasting changes in brain networks that are detectable even in the resting state^5^. For example, Daws et al.^4^ found that psilocybin therapy for depression was associated with decreased brain modularity and increased global integration up to 3 weeks post-treatment, correlating with symptom improvement. Animal research further indicates that this post-treatment period is marked by enhanced neuroplasticity, which is thought to play a crucial role in the therapeutic efficacy of these substances^8,45^. Together, these findings suggest that the period following psychedelic administration represents a window of heightened plasticity in which the brain may be especially receptive to reorganization and integration of new information^46^, particularly if the post-acute period is accompanied by lasting changes in network dynamics.

The dissociation between the two psychedelic groups within the theta band provides evidence that the post-acute theta connectivity effects are 5-HT_2A_ receptor-dependent, consistent with previous work demonstrating that the acute neural effects of typical psychedelics are attributable to 5-HT_2A_ receptor activity^23^, and extends these findings into the post-acute period. It should be noted, however, that the two psychedelic groups differed significantly in days since last use (PSY: 10.43 ± 7.33 vs. MEO: 16.92 ± 7.24, p = .033), and it is possible that the observed theta dissociation partially reflects differential time-dependent recovery rather than pharmacological specificity alone. Still the fact that theta effects did not strongly correlate with days since use, but remained for weeks after the dose contradicts this interpretation.

In the beta frequency band, both psychedelic groups spent a greater proportion of time in a fronto-parietal connectivity state and exhibited longer mean dwell times compared to controls. Beta oscillations are associated with inter-regional communication and cognitive processing^36,47^, and are thought to carry top-down predictions from higher to lower cortical areas^35,38^. The frontal-dominant topography of this state is consistent with changes in prefrontal network-level functions, such as mood-regulation, modulation of sensory processing, and attention^38^. Further, the fact that this was strongest in the right hemisphere is consistent with recent hypotheses, which suggest a lateralized effect of psychedelics which relax a left-to-right hierarchical relationship^48^. The convergence of both psychedelic groups on this effect is notable given their divergent pharmacological profiles and opposing theta patterns, suggesting that this beta effect may not be purely 5-HT_2A_ driven and may instead involve shared downstream mechanisms or 5-HT_1A_ contributions common to both compounds^28,29^.

The REBUS (RElaxed Beliefs Under pSychedelics) model proposes that psychedelics relax the precision weighting of high-level priors, reducing top-down (feedback) constraint and allowing greater bottom-up (feedforward) information flow^46^. Within the predictive coding framework, distinct frequency bands serve distinct roles in cortical information processing. Theta oscillations (5–8 Hz) have been implicated in feedforward information flow across the cortical hierarchy^35–37^, while beta oscillations (15–25 Hz) are associated with top-down predictive signaling, maintaining and communicating predictions from higher to lower cortical areas^35,38^. Increased prominence of a fronto-parietal beta state found here suggests an alteration in this top-down signaling after psychedelics. Likewise, the sustained theta hyper-connectivity observed in the typical psychedelic group may therefore reflect a lasting enhancement of feedforward processing, consistent with the REBUS model’s proposal that psychedelics increase bottom-up drive relative to top-down modulation^46^.

Elevated neural entropy during the acute state is also understood as a corollary of the REBUS hypothesus^19–21,49^, and has been linked to acute revision of maladaptive beliefs^49^. While entropy itself may not persist beyond the acute pharmacological state (see, e.g., figure 1), the well-documented afterglow effects, including changes to mindfulness, mood, and behavioral flexibility lasting up to a month, suggest that lasting alterations in feedforward/feedback network dynamics may emerge when examined at the level of specific frequency bands^16^.

Additionally, 5-HT_2A_ receptor agonism, the primary mechanism of action of typical psychedelics, is thought to weaken top-down modulation while indirectly enhancing bottom-up sensory processing^46^. Another study with this same group of individuals during a predictive saccade paradigm corroborates this effect^50^. Such a shift in hierarchical cortical dynamics could be due to the high concentration of 5-HT_2A_ receptors in cortical layer 5, where top-down connections originate. Activation of these receptors increases cortical excitability and spontaneous activity in layer 5 pyramidal neurons, disrupting normal top-down control^51^. As a consequence of this disrupted top-down modulation, bottom-up sensory processing becomes relatively enhanced^23^. Importantly, 5-HT_2A_ receptors are more prevalent in sensory regions^52^, which may explain the intensified sensory experiences associated with typical psychedelics. In contrast, 5-MeO-DMT has greater affinity for 5-HT_1A_ receptors, which tend to inhibit cortical neuronal activity^23^. 5-HT_1A_ receptors are more concentrated in frontal and temporal lobes^53^, potentially supporting more stable top-down control and contributing to the unique phenomenological effects of 5-MeO-DMT.

The theta connectivity data align with these receptor-level differences. While the typical psychedelic group preferentially visited and lingered in the global hyper-connected theta state, the 5-MeO-DMT group did not exhibit this pattern, instead showing significantly less time in this state compared to the typical psychedelic group. If theta hyper-connectivity reflects enhanced feedforward processing, then the absence of this effect in 5-MeO-DMT users may indicate that feedforward information flow is not similarly enhanced following 5-MeO-DMT use, in contrast to beta-band effects, which may reflect alterations in frontal networks. This finding is also consistent with the phenomenological profile of 5-MeO-DMT, which is characteristically less perceptually driven and more often described as ego dissolution or void-like states^28^, rather than the sensory intensification typical of psilocybin and LSD.

We also identified a group by state effect on alpha connectivity, however the pattern of this effect proved to be complex, as no individual groupwise comparisons for any state proved significant. This suggests that there may be some shifts in alpha networks after a dose. Alpha band is commonly reported as the most robustly altered during acute psychedelic experiences^54,55^. Alpha oscillations serve an inhibitory gating function, regulating the flow of sensory information through thalamocortical circuits^56^, yet also track cortico-cortical circuits as well, often in parallel, involving non-overlapping microcircuitry^57^. Whether shifts in alpha dynamics after a dose reflects altered thalamocortical circuits, corticocortical circuits, or some mixture of both remains to be determined.

Lempel-Ziv Complexity analysis revealed no significant differences between groups, indicating that overall signal complexity had returned to baseline levels in the post-acute period. This normalization suggests that the entropy-generating mechanisms that characterize the acute psychedelic state resolve relatively quickly after a dose, while the lasting effects operate through different neurobiological mechanisms, specifically, frequency specific connectivity changes reported here.

### 4.1 Limitations

Although our findings are intriguing, there are a number of limitations to this study to be addressed in future work. First, there is a gap in the age between groups, with the 5-MeO-DMT group being slightly older (47.17 ± 10.49 versus 37.44 ± 15.08 for controls and 35.71 ± 13.21 for psilocybin/LSD users). This may have had an influence on findings presented in this paper, particularly counteracting any effects of 5-MeO-DMT use on increasing network variability, as measured by number of transitions. However, this age range is still that of stable adulthood and most age effects are fairly minor at this age in the static connectivity literature^58,59^. The effect of age on dFC is an active area of research^60^, and in future work we plan to study the effect of this potential confound and contribute to this growing body of literature. Further, age and sex were both included as a fixed effects in our models, and did not clearly correlate with any primary measure across subjects in the control group.

Additionally, the psilocybin/LSD group had a higher proportion of frequent marijuana users compared to the other two groups, which could confound the results (table 2). As with age, the effects of frequent marijuana use on dFC are not well studied in the literature; however, static FC studies have shown that frequent marijuana use does have complex and lasting impact on global functional connectivity^61^. Thus, future work investigating this potential confound would not only solidify our results here, but contribute to understanding the interaction with frequent marijuana use and dFC. Notably, the two psychedelics groups did not differ in this measure, yet showed divergence in theta connectivity.

The use of dFC provides advantages, as discussed above, but also requires that certain choices be made by researchers, which could influence results. For determining the window size in dFC, we have followed precedent in the literature by selecting a window length of 5 seconds; however, as with the epoch length in static connectivity^62^ it is well established in the EEG literature that the size of a window for sliding window analyses may affect the resulting dynamics observed in connectivity^63^. The obvious next direction for this choice in our work is to systematically study the effect of window size on our analysis. Additionally, a future extension of the methods taken in this work may utilize state of the art approaches for an adaptive window size^64^. In this work, we use K-Means clustering to detect states from wPLI windows. The selection of the number of clusters (K) for K-means clustering, though guided by the elbow criterion, remains a qualitative decision. Future studies can explore alternative clustering methods, such as hierarchical clustering and DBSCAN^65^, which do not require pre-determining the number of clusters. Additionally, some of the assumptions of K-Means such as the dependence on Euclidean distance or assumption of a relatively spherical distribution within clusters may lead to bias in the determination of states. Other clustering methods such as Gaussian Mixture Models may help with this, but still require a predetermination of the number of states like K-Means or the selection of other hyper-parameters which may affect clustering results. Importantly, no comprehensive comparison of different clustering methods has been performed for dynamic functional connectivity in EEG or other modalities. This and other dFC studies utilizing clustering methods would benefit immensely from future work studying the various effects different methods can have on dynamic functional connectivity.

### 4.2 Future Directions

Our study of the effect of psychedelics on patterns in dFC suggests several intriguing paths for future work. While the limitations addressed above suggest obvious paths for studying demographic effects or long-term variations in dFC patterns following psychedelic use, our work provides additional possible extensions in the form of future study designs, methodological extensions to dFC, and comparisons to other connectivity metrics. For example, future research is needed to directly assess the link between dFC characteristics and cognitive flexibility and belief-revision after psychedelic use. Notably, Zeifman et al.^49^ demonstrated that belief revision persists into the afterglow despite neural entropy returning to baseline, leaving open the question of what neural mechanisms facilitate this process. The feedforward/feedback dynamics observed here represent one candidate worthy of direct investigation. Additionally future work should aim to address the relationship between the observed measures of increased entropy during the acute treatment period and the long-term effects on dFC observed here. These experiments could provide empirical evidence to support a largely theoretical framework explaining psychedelics’ effects on neural activity and how these effects lead to increases in cognitive flexibility and the reconfiguration of pathological beliefs. Furthermore, our finding that theta and beta connectivity effects dissociate along pharmacological lines raises the question of whether the feedforward and feedback components of the predictive coding hierarchy are independently modifiable, and whether these distinct neural profiles correspond to different therapeutic mechanisms across psychedelic compounds.

### 4.3 Conclusion

This study utilized dynamic functional connectivity to characterize distinct post-acute neural signatures of typical and atypical psychedelic use. Notably, a hallmark effect seen in the acute psychedelic state – elevated entropy – did not persist into the post-acute period. What did persist is a reorganization of connectivity dynamics within the feedforward and feedback channels of the cortical hierarchy. In the theta band, only typical psychedelic users showed increased occupancy of a global hyper-connectivity state, consistent with enhanced feedforward processing mediated by 5-HT_2A_ receptor agonism. In the beta band, both psychedelic groups preferentially occupied a frontal-parietal dominant connectivity state, suggesting a shared alteration of feedback-associated dynamics that may not be exclusively 5-HT_2A_ dependent. These findings suggest that the lasting neural impact of psychedelics is not sustained entropy but shifts in cortical networks, and that typical and atypical psychedelics may engage partially dissociable pathways in this process.

