## Supplemental Figures 1-12 for "A dominant frontoparietal beta oscillatory brain state in the days after psilocybin and 5-MeO-DMT"

Supplemental materials for “A dominant fronto-parietal beta oscillatory brain state in the days after psilocybin and 5-MeO-DMT”

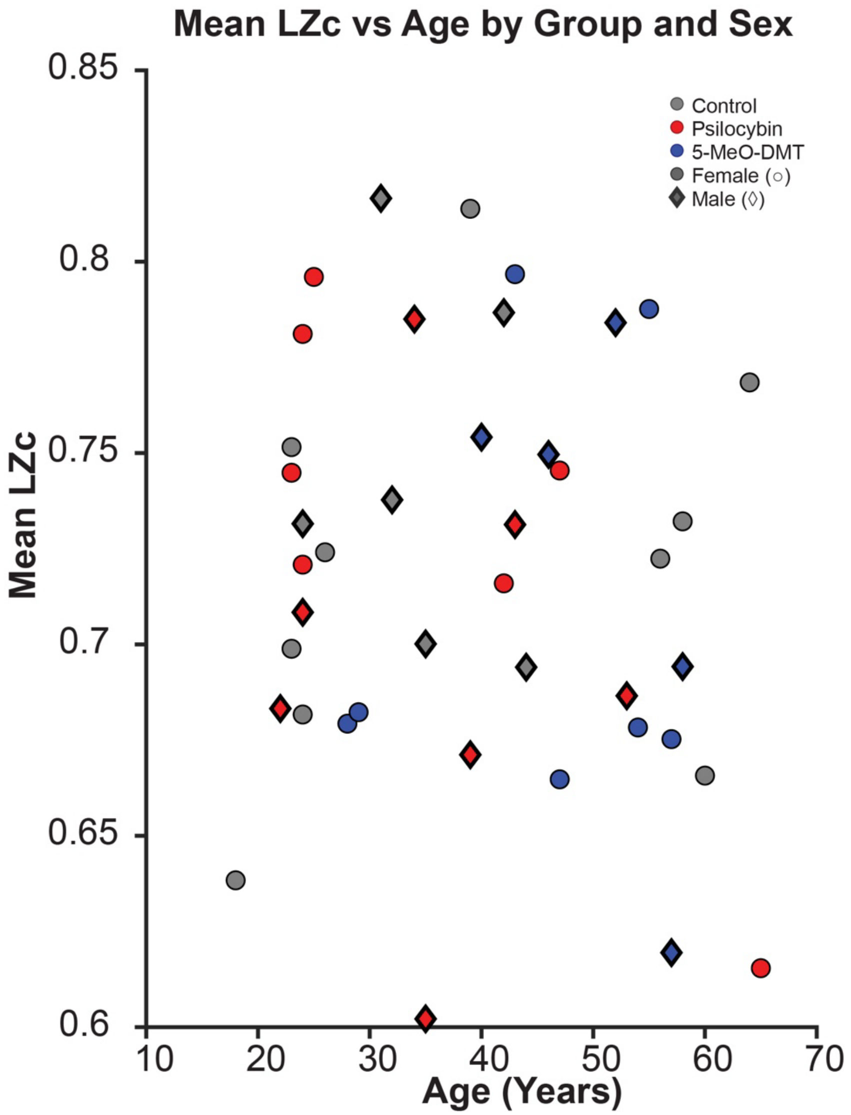


**Supplemental Figure S1.** Scatter plot of average global lempel-ziv complexity (across scalp) vs age of subjects. Males are diamonds.

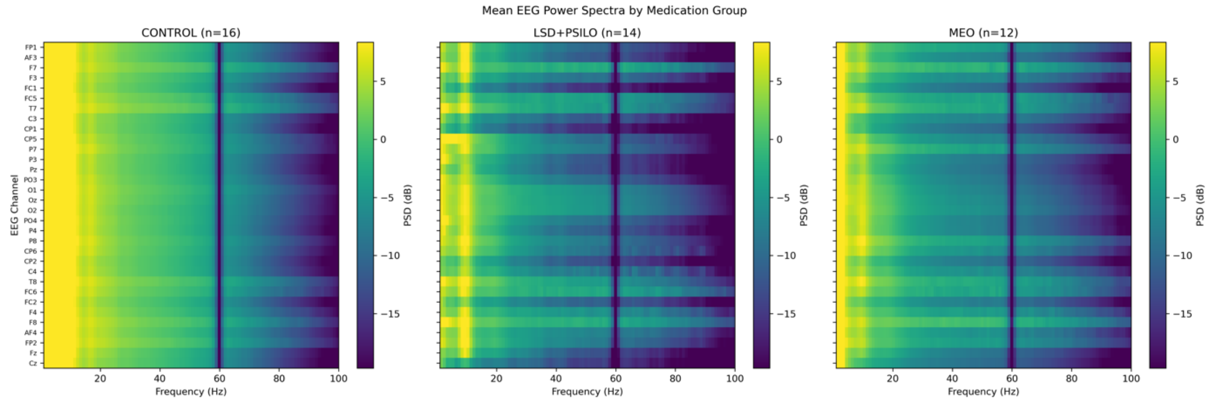


**Supplemental figure S2:** Average resting power for each electrode (y-axis) for each frequency (x-axis) averaged over subjects within each group, over the full 5-minute resting interval (after artifact removal).

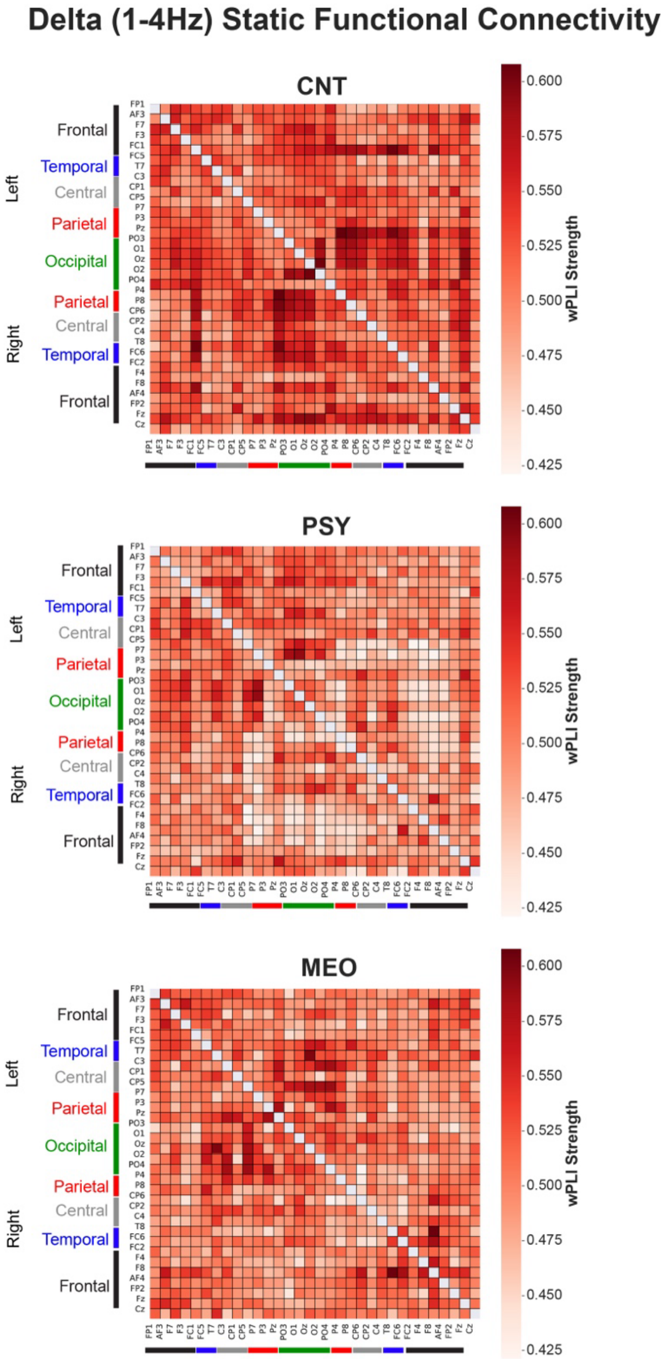


**Supplemental figure S3: Delta-band.** Functional connectivity matrices (weighted phase-locking index) across electrodes, averaged within groups over the full resting period, prior to state segregation.

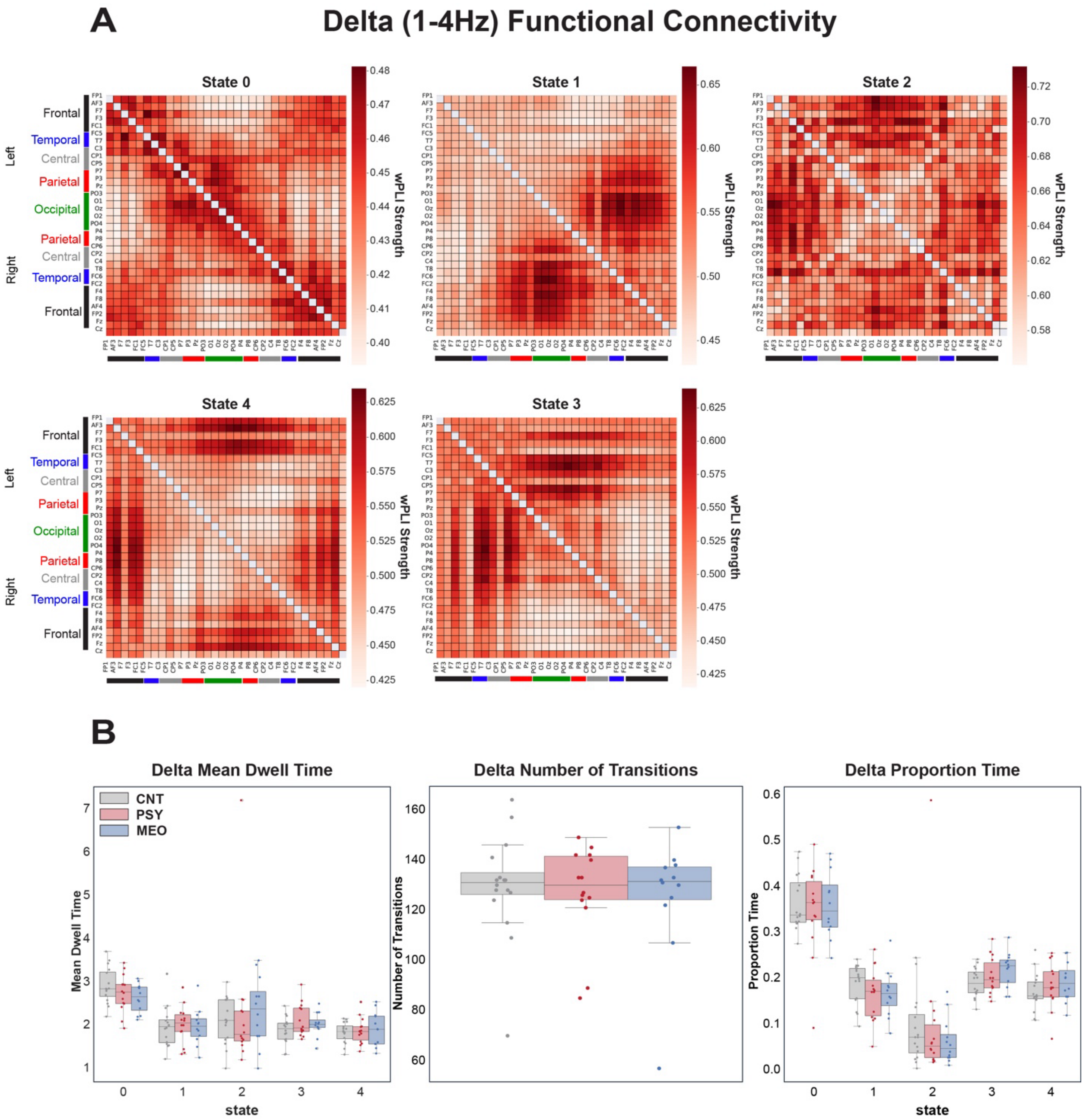


**Supplemental figure S4: Delta-band.** A) Functional connectivity matrices (weighted phase-locking index), plotted across electrodes, averaged over all subjects for each of 5 states. B) Box plots (median and IQR), with individual subject values overlaid (dots) for each group, depicting mean dwell time in each state, number of transitions between states, and overall proportion of time spent within each state, averaged over the entire 5-minute rest period.

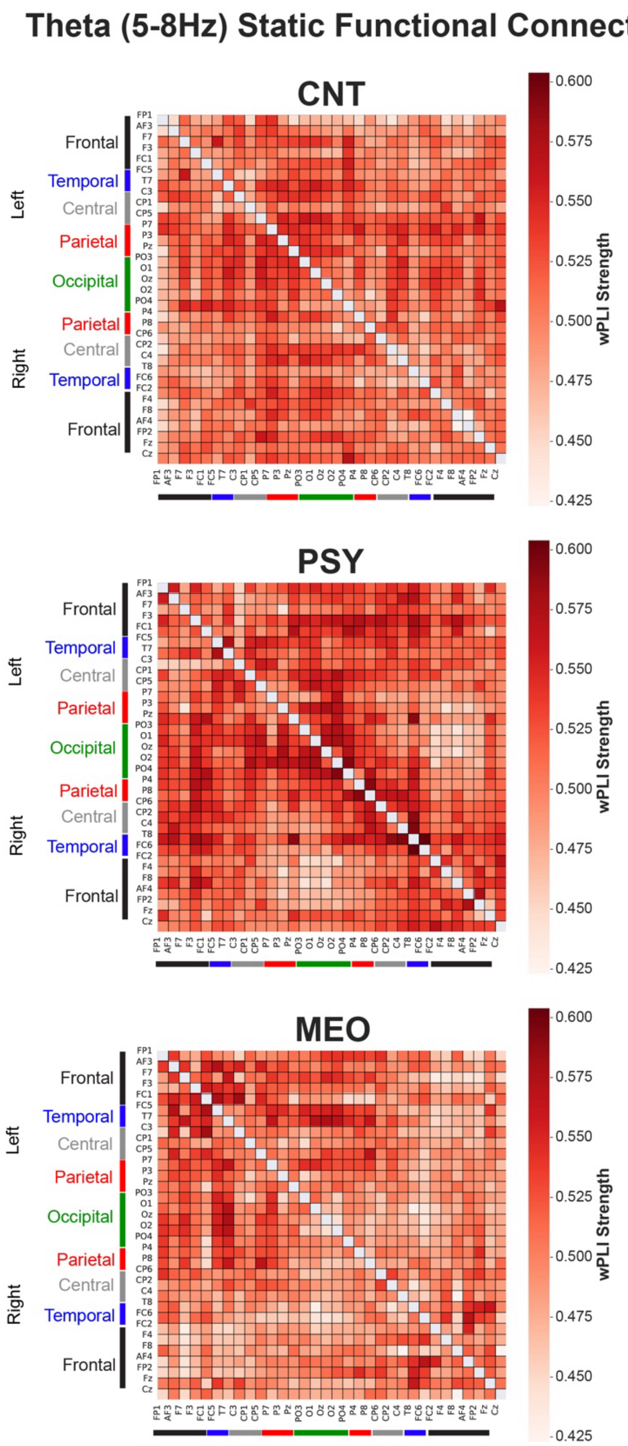


**Supplemental figure S5: Theta-band.** Functional connectivity matrices (weighted phase-locking index) across electrodes, averaged within groups over the full resting period, prior to state segregation.

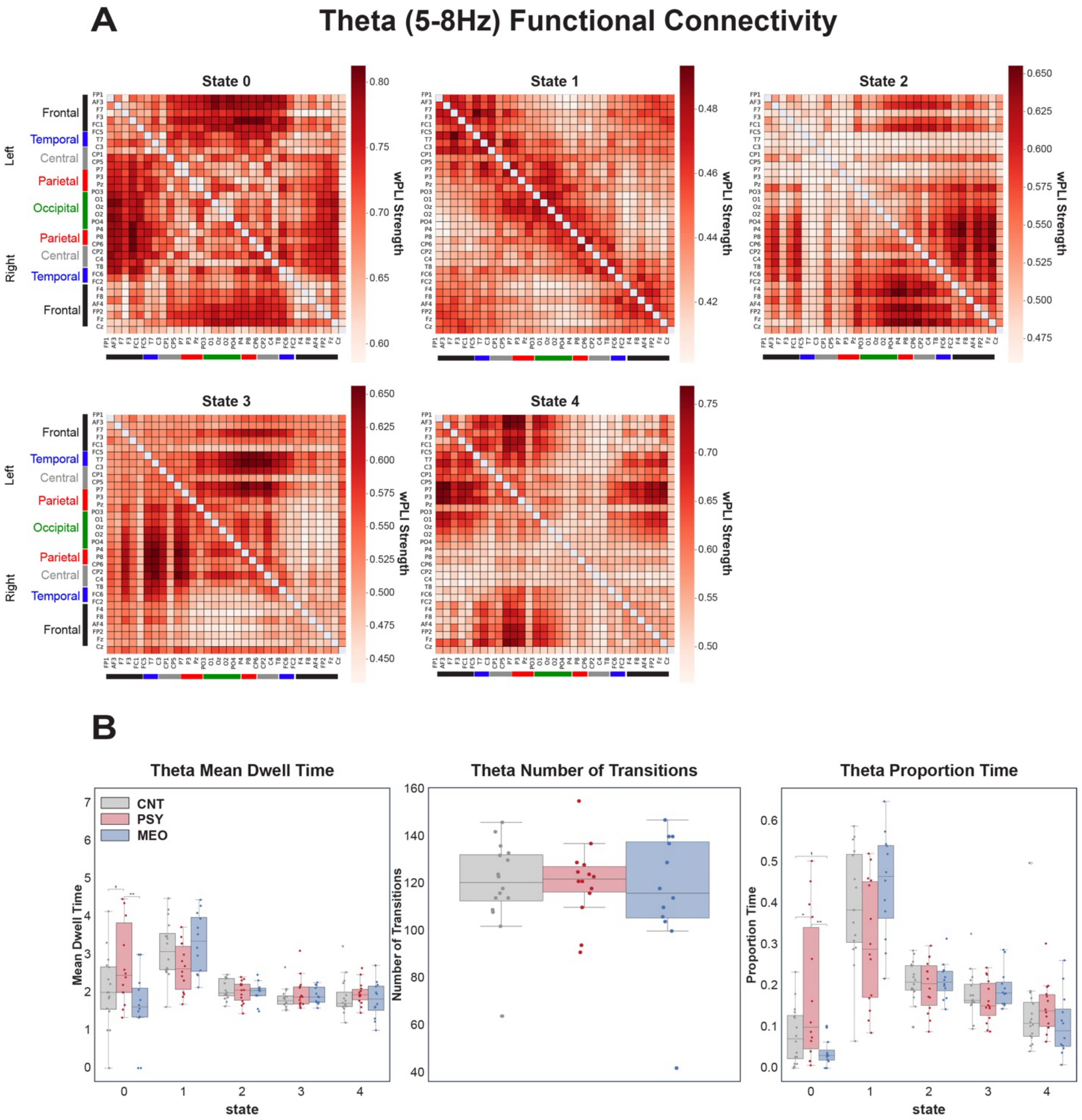


**Supplemental figure S6: Theta-band.** A) Functional connectivity matrices (weighted phase-locking index), plotted across electrodes, averaged over all subjects for each of 5 states. B) Box plots (median and IQR), with individual subject values overlaid (dots) for each group, depicting mean dwell time in each state, number of transitions between states, and overall proportion of time spent within each state, averaged over the entire 5-minute rest period.

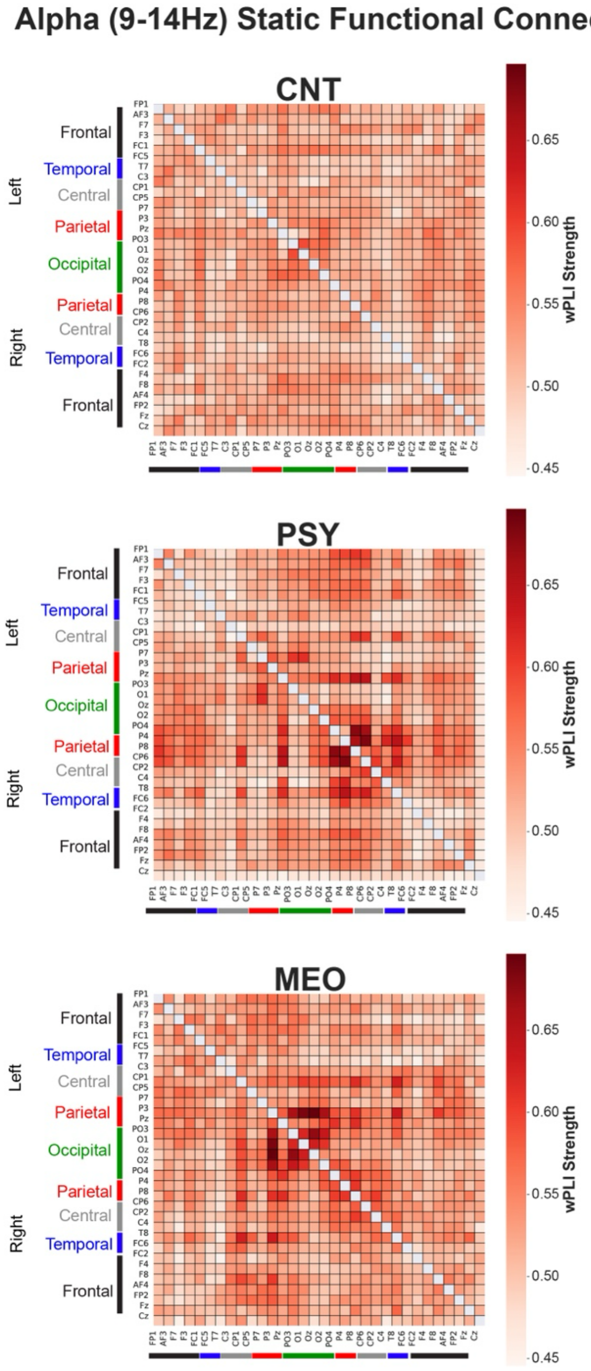


**Supplemental figure S7: Alpha-band.** Functional connectivity matrices (weighted phase-locking index) across electrodes, averaged within groups over the full resting period, prior to state segregation.

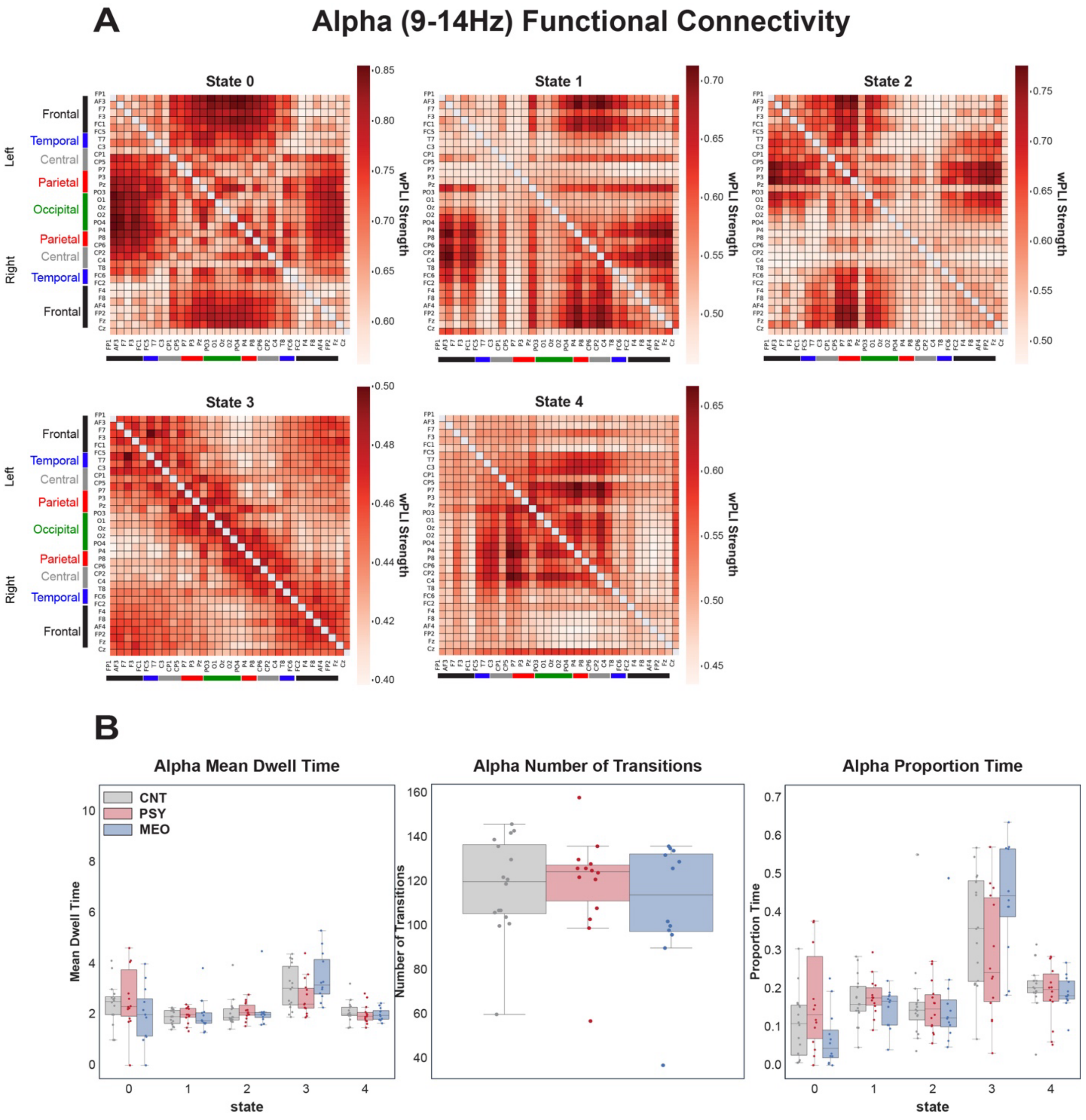


**Supplemental figure S8: Alpha-band.** A) Functional connectivity matrices (weighted phase-locking index), plotted across electrodes, averaged over all subjects for each of 5 states. B) Box plots (median and IQR), with individual subject values overlaid (dots) for each group, depicting mean dwell time in each state, number of transitions between states, and overall proportion of time spent within each state, averaged over the entire 5-minute rest period.

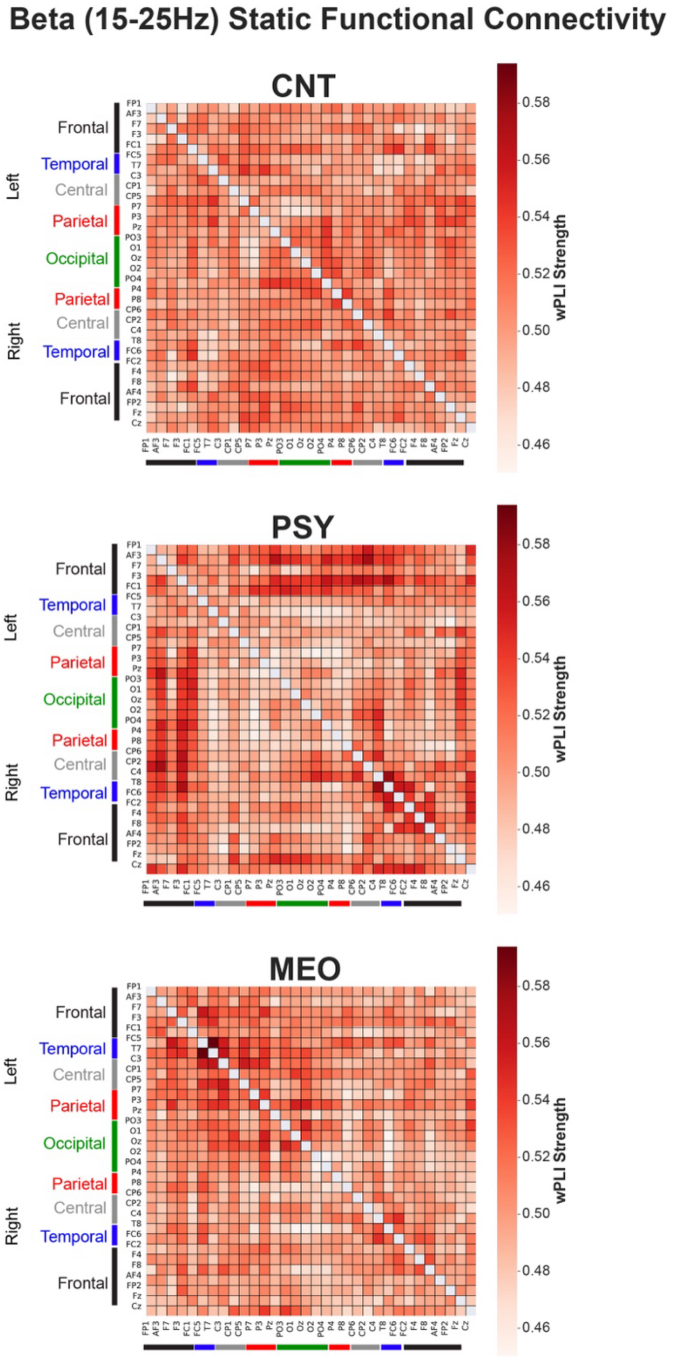


**Supplemental figure S9: Beta-band.** Functional connectivity matrices (weighted phase-locking index) across electrodes, averaged within groups over the full resting period, prior to state segregation.

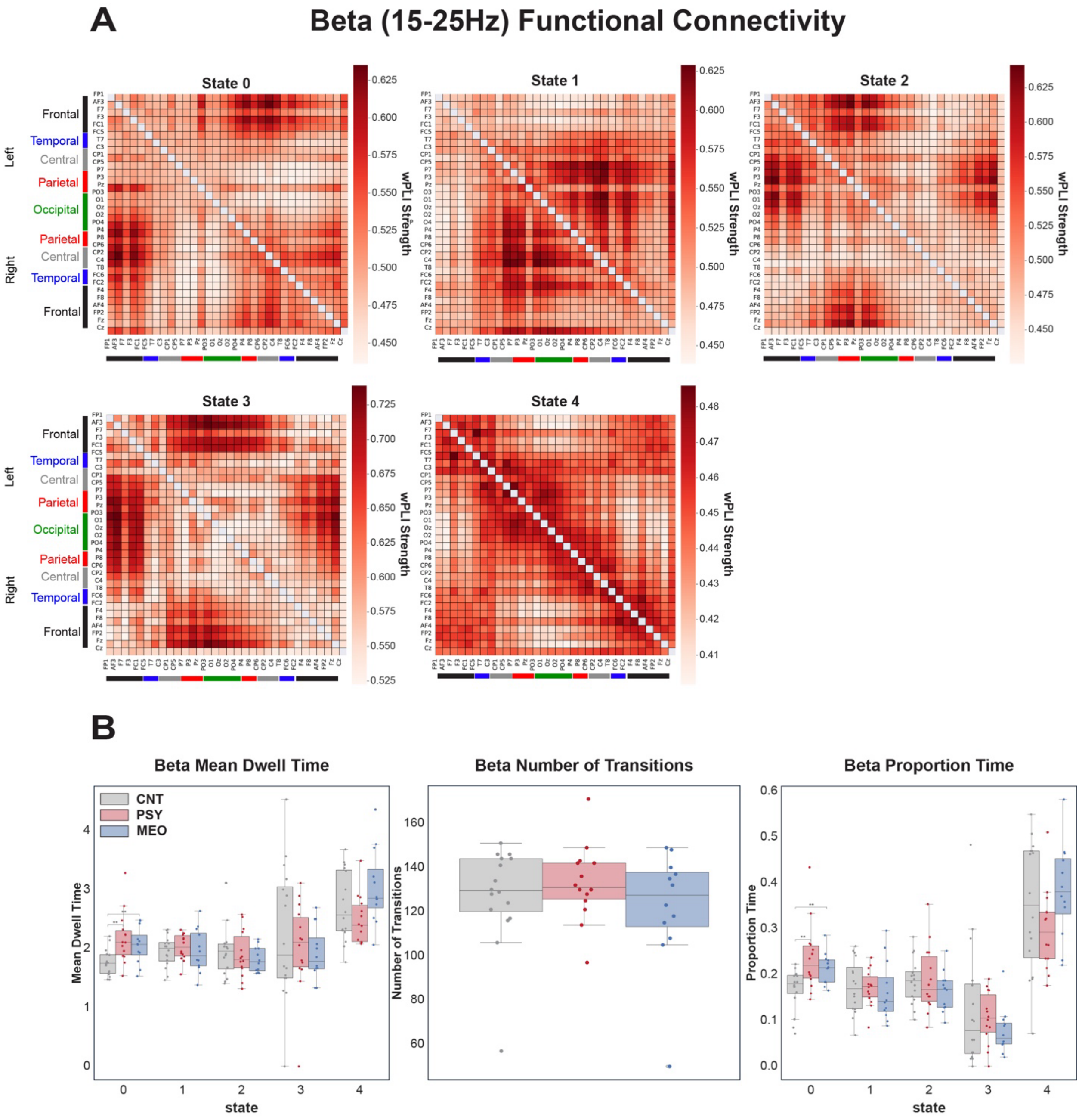


**Supplemental figure S10: Beta-band.** A) Functional connectivity matrices (weighted phase-locking index), plotted across electrodes, averaged over all subjects for each of 5 states. B) Box plots (median and IQR), with individual subject values overlaid (dots) for each group, depicting mean dwell time in each state, number of transitions between states, and overall proportion of time spent within each state, averaged over the entire 5-minute rest period.

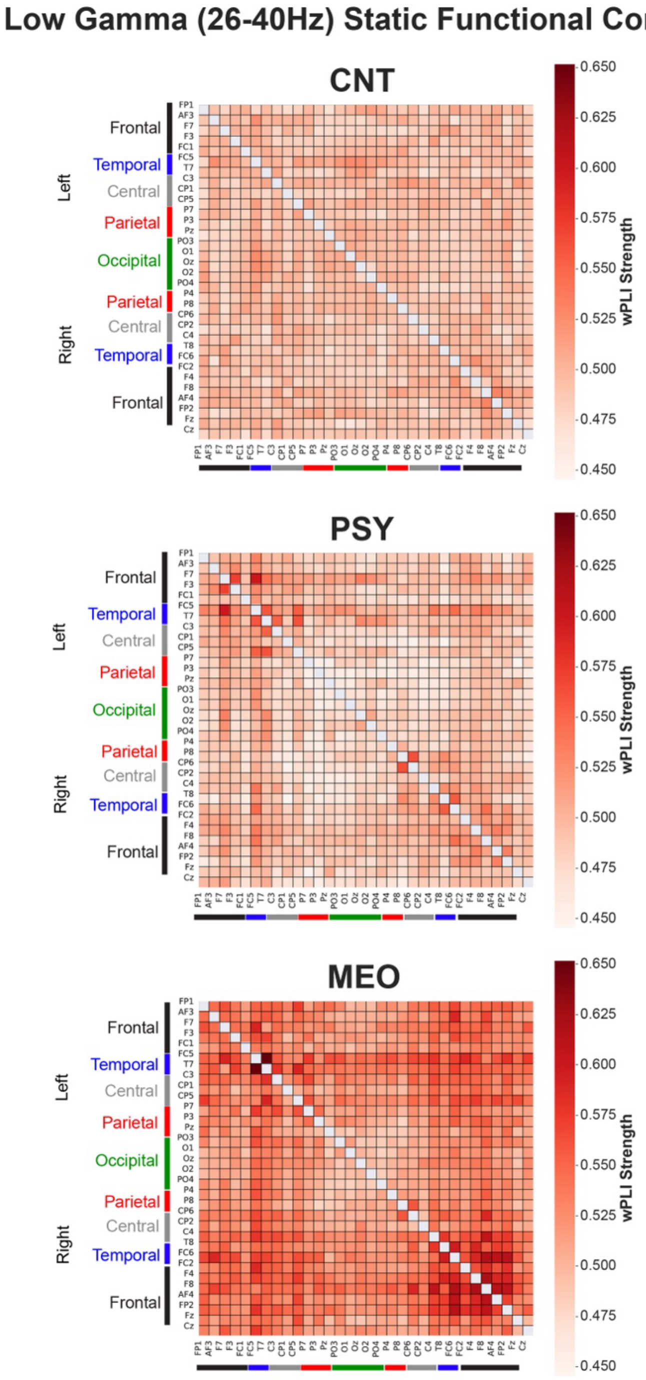


**Supplemental figure S11: Gamma-band.** Functional connectivity matrices (weighted phase-locking index) across electrodes, averaged within groups over the full resting period, prior to state segregation.

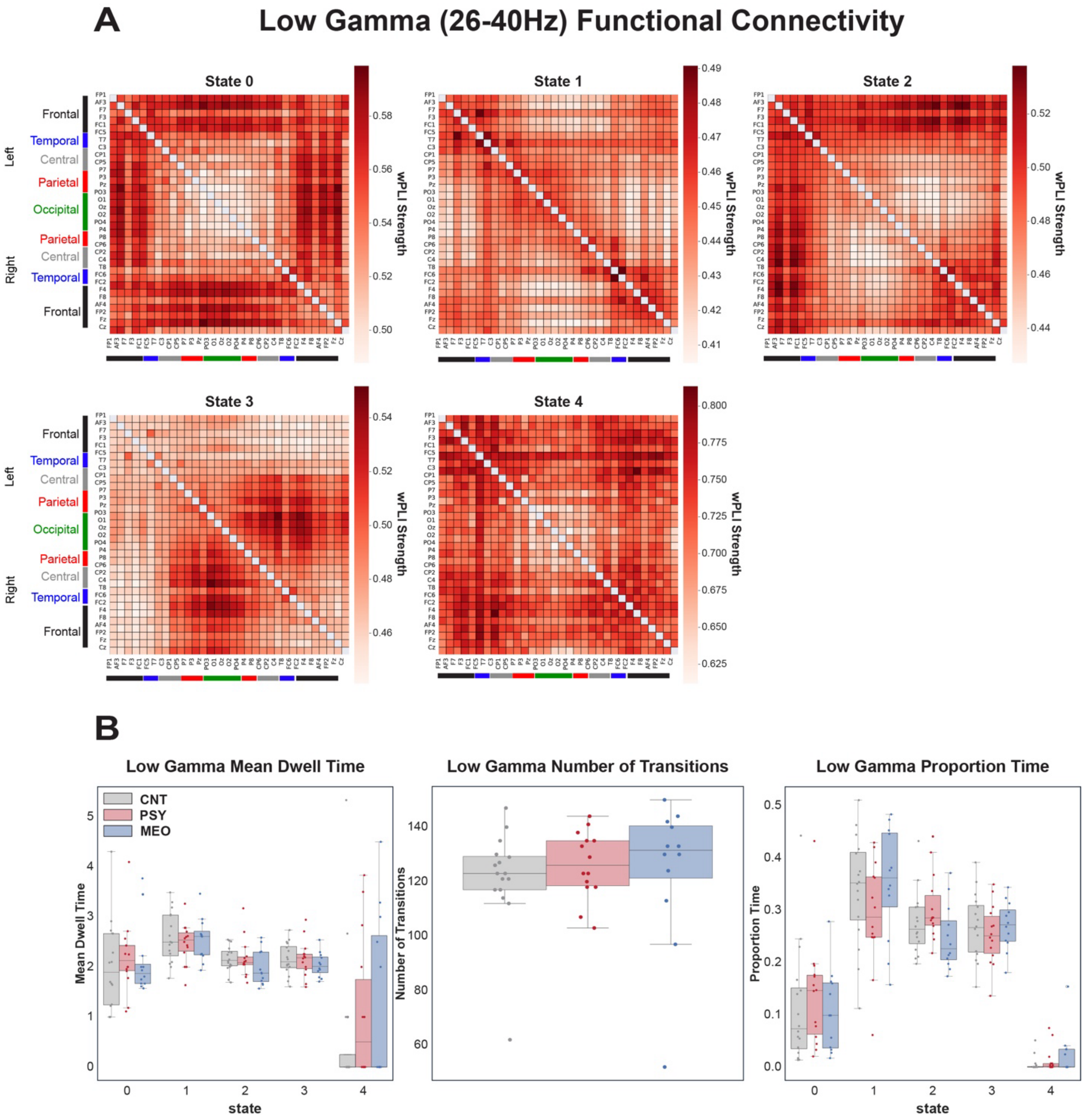


**Supplemental figure S12: Gamma-band.** A) Functional connectivity matrices (weighted phase-locking index), plotted across electrodes, averaged over all subjects for each of 5 states. B) Box plots (median and IQR), with individual subject values overlaid (dots) for each group, depicting mean dwell time in each state, number of transitions between states, and overall proportion of time spent within each state, averaged over the entire 5-minute rest period.
